# Shape-invariant perceptual encoding of dynamic facial expressions across species

**DOI:** 10.1101/2020.07.10.195941

**Authors:** N. Taubert, M. Stettler, R. Siebert, S. Spadacenta, L. Sting, P. Dicke, P. Thier, Martin A. Giese

**Author notes:** **Materials & Correspondence**, Martin Giese. equal contributions (N. Taubert, M. Stettler). equal contributions (P. Thier, Martin A. Giese).

## Abstract

Dynamic facial expressions are crucial for communication in primates. Due to the difficulty to control shape and dynamics of facial expressions across species, it is unknown how species-specific facial expressions are perceptually encoded and interact with the representation of facial shape. While popular neural-network theories predict a joint encoding of facial shape and dynamics, the neuromuscular control of faces evolved more slowly than facial shape, suggesting a separate encoding. To investigate this hypothesis, we developed photo-realistic human and monkey heads that were animated with motion-capture data from monkeys and human. Exact control of expression dynamics was accomplished by a Bayesian machine-learning technique. Consistent with our hypothesis, we found that human observers learned cross-species expressions very quickly, where face dynamics was represented independently of facial shape. This result supports the co-evolution of the visual processing and motor-control of facial expressions, while it challenges popular neural-network theories of dynamic expression-recognition.

## Introduction

Facial expressions are crucial for social communication of human as well as non-human primates^1–4^, and humans can learn facial expressions even of other species^5^. While facial expressions in everyday life are dynamic, specifically expression recognition across different species has been studied mainly using static pictures of faces^6–10^. A few studies have compared the perception of human and monkey expressions using movie stimuli, finding overlaps in the brain activation patterns induced by within- and cross-species expression observation in humans as well as in monkeys^11,12^. Since natural video stimuli provide no accurate control of the dynamics and form features of facial expressions, it is unknown how expression dynamics is perceptually encoded across different primate species, and how it interacts with the representation of facial shape.

In primate phylogenesis the visual processing of dynamic facial expressions has co-evolved with the neuromuscular control of faces^13^. Remarkably, the structure and arrangement of facial muscles is highly similar across different primate species^14,15^, while face shapes differ considerably, e.g. between humans, apes, or monkeys. This motivates the following two hypotheses: 1) The phylogenetic continuity in motor control should facilitate fast learning of dynamic expressions across primate species; and 2) the different speeds of the phylogenetic development of the facial shape and its motor control should potentially imply a separate visual encoding of expression dynamics and basic face shape.

We investigated these hypotheses, exploiting advanced methods from computer animation and machine learning, combined with motion capture in monkeys and humans. We designed highly-realistic three-dimensional human and monkey avatar heads by combining structural information derived from 3D scans, multi-layer texture models for the reflectance properties of the skin, and hair animation. Expression dynamics was derived from motion capture recordings on monkeys and humans, exploiting a hierarchical generative Bayesian model to generate a continuous motion-style space. This space includes continuous interpolations between two expression types (‘anger’ vs. ‘fear’), and human- and monkey-specific motion. Human observers categorized these dynamic expressions, presented on the human or the monkey head model, in terms of the perceived expression type and species-specificity of the motion (human vs. monkey expression).

Consistent with our hypotheses, we found very fast cross-species learning of expression dynamics with a more precise tuning for human-compared to monkey-specific expressions. Most importantly, the perceptual representation of expression dynamics was largely independent of the facial shape (human vs. monkey). Perceptual responses were determined by the coordinates of the stimuli in the motion style space, and did not depend on the matching of face species with the species-specificity of the motion. Our results were highly robust against substantial variations in the expressive stimulus features. They specify fundamental constraints for the computational neural mechanisms of dynamic face processing and challenge popular neural network models, accounting for expression recognition by the learning of sequences of key shapes^4^.

## Results

Exploiting photo-realistic human and monkey face avatars, we investigated the perceptual representations of dynamic human and monkey facial expressions in human observers. The dynamic avatars were created by combining advanced computer animation methods with motion capture in both primate species (Figures 1A and 1B).

**Figure 1.**
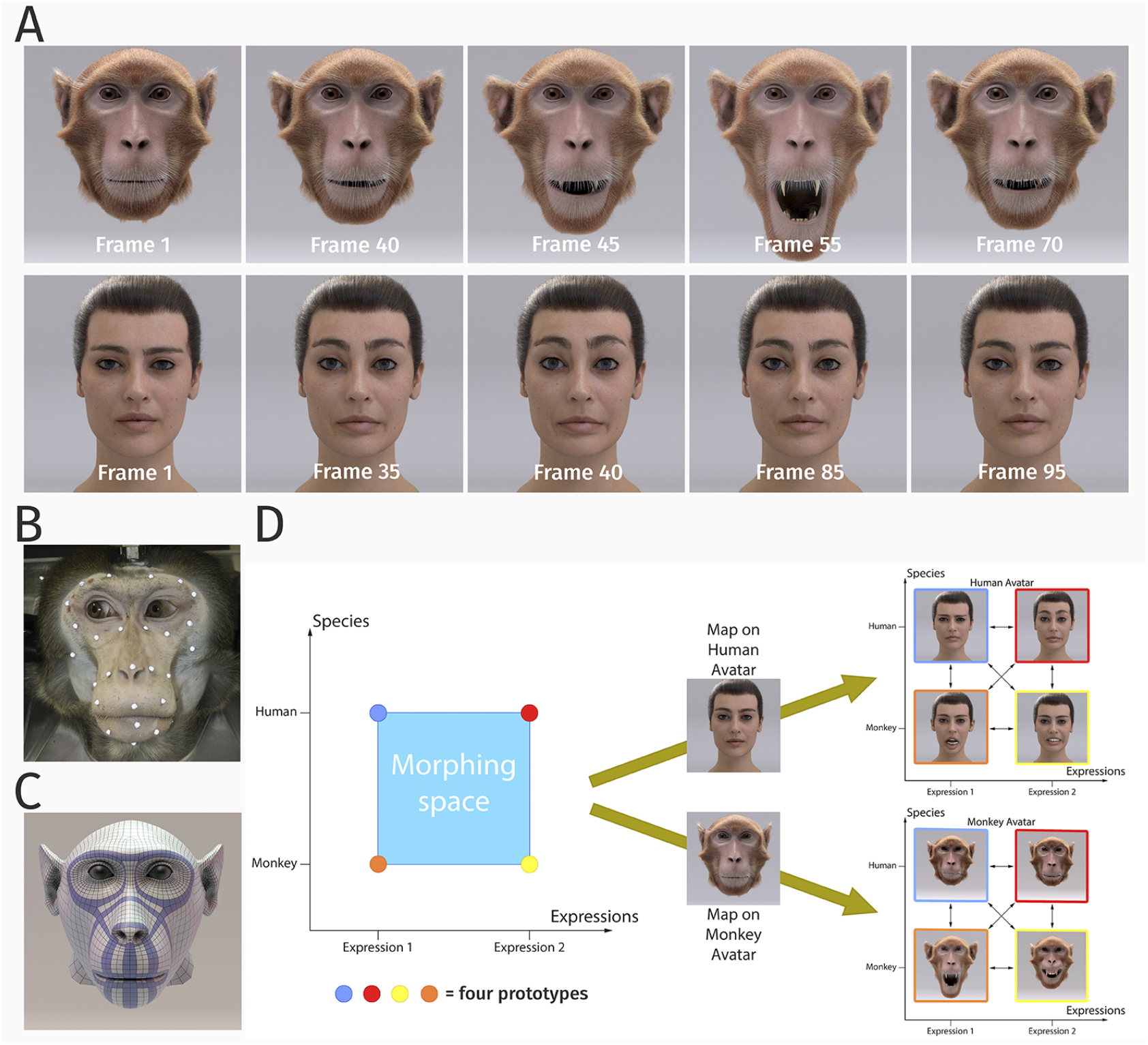
Stimulus generation and paradigm. (A) Frame sequence of a monkey and a human facial expression. (B) Monkey motion capture with 43 reflecting facial markers. (C) Regularized face mesh whose deformation is controlled by an embedded elastic ribbon-like control structure that is optimized for animation. (D) Stimulus set. We generated 25 motion patterns, spanning up a two-dimensional style space with the dimensions ‘expression’ and ‘species’ by interpolation between two expressions (‘anger’ and ‘fear’) and the two species (‘monkey’ and ‘human’). Each motion pattern was used to animate a monkey and a human avatar model.

### Highly realistic dynamic face avatars

We developed a photo-realistic monkey head model, whose degree of realism exceeds the one of all avatars used previously in perception and physiological research^16–18^. It was derived from a structural magnetic resonance scan of a rhesus monkey. The surface of the face was modeled by an elastic mesh structure (Fig 1C) which imitates the deformations induced by the major face muscles of macaque monkeys^15^. The motion of this mesh was specified by motion capture of 43 reflecting markers. Skin surface and fur were modeled in very much detail in order to achieve a high level of realism (Fig. 1A). A similar highly-realistic human avatar model was created based on a commercially available scan-based human face model. Its animation was based on blend shapes, exploiting a multi-channel texture simulation software. Mesh deformations compatible with the human face muscle structure were computed from motion capture data in the same way as for the monkey face (cf. Supplementary Information for details).

The facial motion of the avatars was based on motion capture data from humans and monkeys. We recorded two expressions (prototypes), *anger/threat* and *fear* from both species. Facial movements of humans and monkeys are quite different^14^, so that our participants, who all had no prior experience with macaque monkeys, needed to be familiarized briefly with the monkey expressions. In order to study the structure of the perceptual representation parametrically, we generated a continuous dynamic expression space by morphing between four prototypical expressions, ‘anger/threat’ and ‘fear’, each executed by humans and monkeys. Interpolated patterns were generated by a Bayesian generative model that was trained with examples of the four prototypical face movements, resulting in a style space that included a total of 25 facial movements that interpolate between the prototypes (see Supplementary Information for details on the algorithm). Each generated motion pattern can be parameterized by a two-dimensional style vector (*e*, *s*), where the first component *e* specifies the expression type (e = 0: expression 1 (‘fear’), and *e* = 1: expression 2 (‘anger/threat’)), and where the second variable *s* the species-specificity of the motion (s = 0: monkey, and s = 1: human). The resulting patterns corresponded to equidistant points between 0 and 1 along these two style axes (Figure 1D). The 25 generated facial movements were presented on the monkey as well as on the human avatar in order to study how the basic shape of the avatar influences the perception of the dynamic facial expressions. A control experiment (see Supplementary Information) verifies that faces animated with the motion morphs are not perceived as less natural than faces animated with original motion capture data.

### Dynamic expression perception is largely independent of facial shape

In our first experiment, we used the original dynamic expressions of humans and monkeys as prototypes and presented morphs between them, separately, on the human and the monkey avatar face. Prior to the experiment, participants were familiarized with the prototype stimuli, repeating each stimulus at maximum 10 times and stopping as soon as the prototypes were recognized reliably. Motions were presented in a randomized order, and in separate blocks for the two avatars. The expression movies had a duration of 5 s and showed the face going from a neutral expression to the extreme expression, and back to neutral (Fig. 1A). Participants observed 10 repetitions of each stimulus in block-randomized order. They had to decide whether the observed stimulus was looking more like a human or a monkey expression (independent of the avatar type), and whether the expression was rather ‘anger/threat’ or ‘fear’. The resulting two binary responses in each trial can be interpreted as assignment of one out of four classes to the stimulus (expression 1 vs. 2, either monkey- or human-specific movement).

In order to model these categorization results as a function of the position of the stimulus in the two-dimensional motion style space, we approximated the classification probabilities of the four classes by a logistic multinomial regression model. The resulting fits are shown in Figures 2A and 2B for the two avatar types. The class probabilities *P_i_* for the four classes were approximated by a Generalized Linear Model of the form:

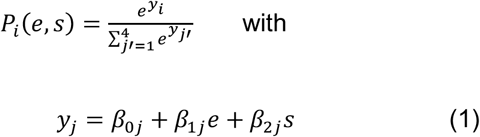

where *P_i_* is the probability of class *i* as a function of the position of the stimulus in morphing space. We tested also further variants of linear models for which the prediction *y_i_* depended on more or less variables as predictors. A comparison of the prediction accuracies of these models is shown in Figure 3A for the monkey avatar, where results for the human avatar are very similar. Model comparison exploiting the Bayesian Information Criterion shows that (1) is the most compact model that explains the classification data with high accuracy. Specifically, models only including the predictors *e* or *s* provided significantly worse fits, and a model with an additional predictor of the form *e* * *s* did not result in better predictions. Likewise, models that contained the average amount of optic flow as additional predictor did not result in higher accuracy (see Table 1). These results imply an almost entirely linear dependence of the classification model (1) on the style space coordinates (*e*, *s*). Consequently, we used this model as basis for our further analyses.

**Figure 2.**
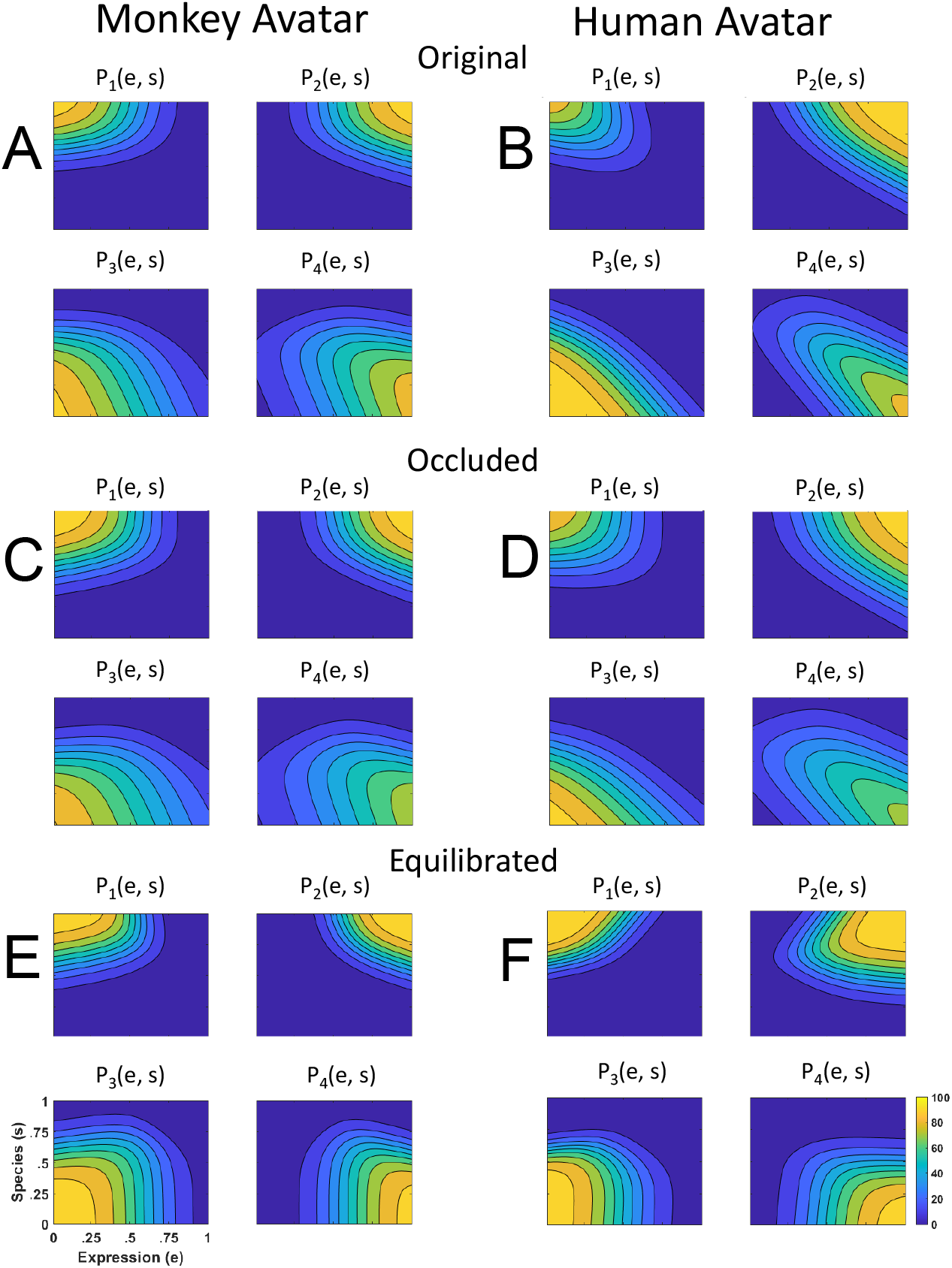
Discriminant functions *P_i_*(*e*, *s*) fitted to the classification responses. Classes correspond to the four prototype motions, as specified in Fig. 1D (*i* = 1, 2: monkey, and *i* = 3, 4: human motion). (A) Discriminant functions for the stimulus set created using original motion-captured expressions of humans and monkeys as prototypes, for presentation on a monkey and a human avatar. (B) Same results for stimuli with occluded ears. (C) Results for a stimulus set derived from prototypes that were equilibrated with respect to the amount of local motion or deformation information.

**Figure 3.**
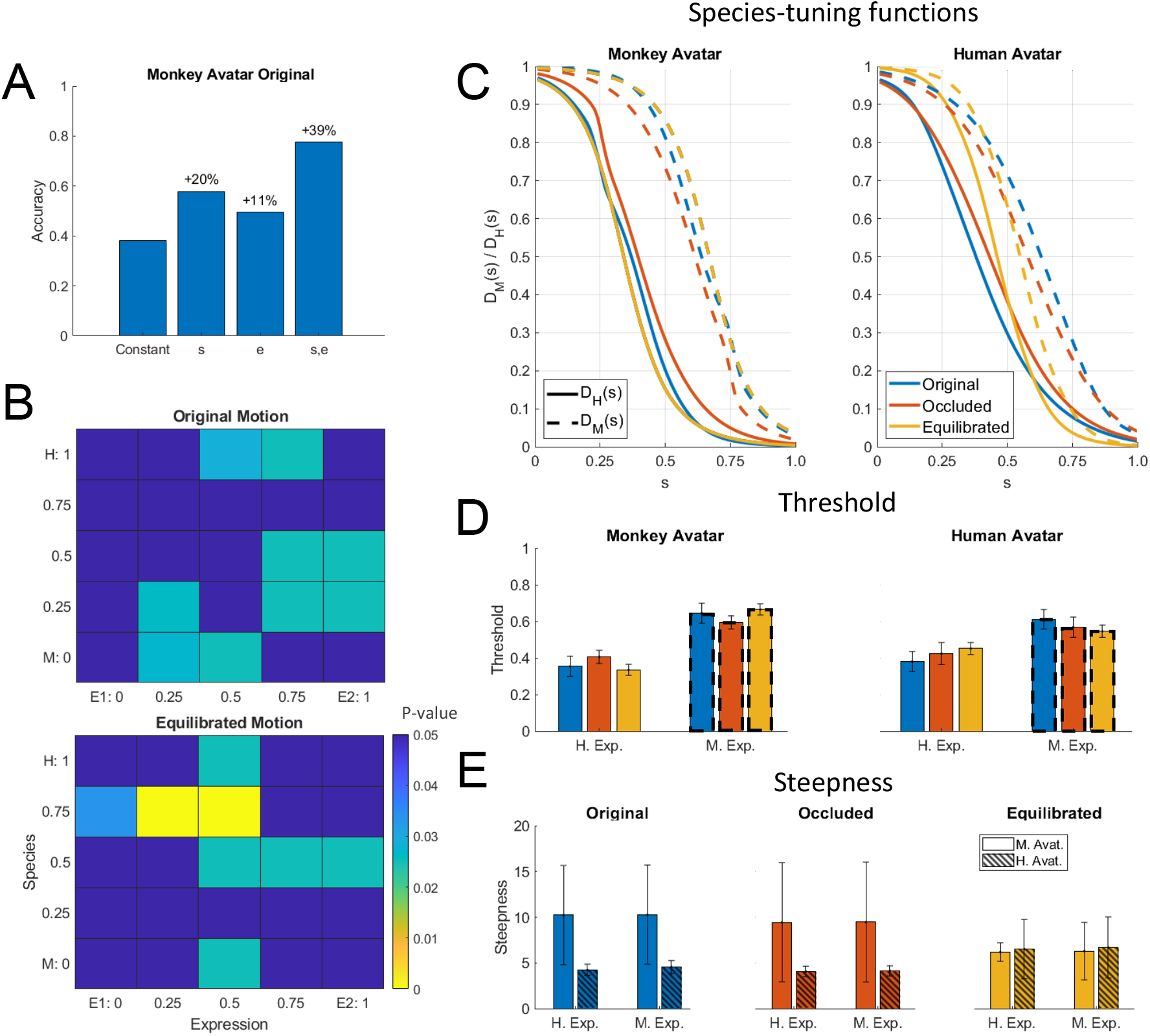
Statistical analysis of the results. (A) Accuracy of the fits of the discriminant functions using Generalized Linear Models (GLMs) with different sets of predictors. Numbers indicate change in accuracy compared to the constant model. (B) Significance levels (Bonferroni-corrected) of the differences between the multinomially distributed classification responses for the 25 motion patterns, presented on the monkey and human avatar. (C) Fitted tuning functions *D*_H_(*s*) (solid lines) and *D*_M_(*s*) (dashed lines) for the categorization of patterns as monkey vs. human expressions, separately for the two avatar types. Different line styles indicate the experiments using original motion captured motion, stimuli with occluded ears, and the experiment using prototype motions that was equilibrated for the amount of motion / deformation across prototypes. (D) Thresholds of the tuning functions for the three experiments for presentation on the human and monkey avatar. (E) Steepness of the tuning functions at the threshold points for the experiments with and without equilibration of the prototype motions (and without occlusions). (Uniformly colored bars indicate the results for the monkey avatar and dashed bars the ones for the human avatar.)

**Table 1.** Model Comparison. Results of the Accuracy and the Bayesian Information Criterion (BIC) for the different logistic multinomial regression models for the stimuli derived from the original motion (no occlusions) for the monkey and the human avatar. The models included the following predictors: Model 1: constant; Model 2: constant, *s*; Model 3: constant, *e*; Model 4: constant, *s*, *e*; Model 5: constant, *s*, *e*, product *s*·*e* Model 5: constant, *s*, *e*, Optic Flow.

| Model Comparison |  |  |  |  |  |  |  |  |
| --- | --- | --- | --- | --- | --- | --- | --- | --- |
| Monkey Avatar | Model | Accuracy [%] | Accuracy increase [%] | BIC | Parameters | df | $\chi^2$ | p |
|  | Model 1 | 38,29 |  | 7487 | 33 |  |  |  |
|  | Model 2 | 57,86 | 19,56 (rel. to Model 1) | 5076 | 36 | 3 | 2411 | <0,0001 |
|  | Model 3 | 49,49 | 11,2 (rel. to Model 1) | 6125 | 36 | 3 | 1362 | <0,0001 |
|  | Model 4 | 77,53 | 19.7 (rel. to Model 2) | 3586 | 39 | 3 | 1490 | <0,0001 |
|  | Model 5 | 77,53 | 0 (rel. to Model 4) | 3598 | 42 | 3 | -11,997 | 1 |
|  | Model 6 | 77,42 | -0.11 (rel. to Model 4) | 3580 | 42 | 3 | 5,675 | 0,129 |
| Human Avatar |  |  |  |  |  |  |  |  |
|  | Model 1 | 36,84 |  | 7481 | 33 |  |  |  |
|  | Model 2 | 54,22 | 17,38 (rel. to Model 1) | 5541 | 36 | 3 | 1940 | <0,0001 |
|  | Model 3 | 53,56 | 16,72 (rel. to Model 1) | 5847 | 36 | 3 | 1633 | <0,0001 |
|  | Model 4 | 81,56 | 27,35 (rel. to Model 2) | 3420 | 39 | 3 | 2120 | <0,0001 |
|  | Model 5 | 81,35 | -0,22 (rel. to Model 4) | 3309 | 42 | 3 | 112 | <0,0001 |
|  | Model 6 | 81,38 | -0.18 (rel. to Model 4) | 3389 | 42 | 3 | 31,66 | <0,0001 |

The functional forms of the discriminant functions for the human and the monkey avatar (Figure 2 A and B) were very similar. This is confirmed by the fact that the fraction of the variance that is different between these functions divided by the one that is shared does not exceed 10% (*q* = 6.35 %; see Methods). Also, a comparison of the multinomially distributed classification responses between the two avatar types, separately for the different points in morphing space and across participants, revealed no significant differences across all tested points in morphing space (*p* = 0.02, Bonferroni-corrected). Differences tended to be larger especially for intermediate values of the coordinates *e* and *s*, thus for the stimuli with high perceptual ambiguity (Fig. 3B). This result implies that the facial motion of human and monkey facial expressions is encoded largely independently of the basic shape of the avatar (human or monkey). This independence might also explain why many of our subjects were able to recognize *human* facial expressions on the monkey avatar face spontaneously, even without familiarization.

### Tuning is narrower for human-specific than for monkey-specific dynamic expressions

A biologically important question is whether expressions of the own species are processed differently from those of other primate species, potentially supporting an *own-species advantage* in the processing of dynamic facial expressions^19^. In order to characterize the tuning of the perceptual representation for monkey vs. human expressions, we computed tuning functions, marginalizing the discriminant functions belonging to the same species category (*P*_1_ and *P*_2_ belonging to the human, and *P*_3_ and *P*_4_ to the monkey expressions) over the expression dimension *e*. This defines the function 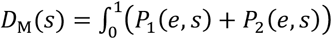 d*e* that characterizes the tuning to monkey expressions as function of the species dimension *s*, and the function 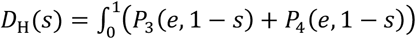 d*e*, which characterizes the tuning to human expressions. In the function *D*_H_(*s*) we flipped the *s*-axis so that the category center also appears for *s* = 0, just as for the function *D*_M_(*s*). Figure 3C shows these two species-tuning functions, revealing smaller tuning width for the human than for the monkey expressions. This observation is statistically confirmed by fitting of the tuning functions by a sigmoidal threshold function. The fitted threshold values *s*_th_ with *D_M_*(*s_th_*), *D_H_*(*s_th_*) = 0.5 are shown in (Fig. 3D). They are significantly smaller for the human expression tuning functions *D*_H_(*s*) than for the monkey expression tuning functions *D*_M_(*s*) for both avatars. This is confirmed by two separate ANOVAs for the two avatar types. These 2-way mixed-model ANOVAs include the expression type (human vs. monkey motion) as within-subject factor, and the stimulus type (original motion, stimuli with occluded ears, or animated with equilibrated motion; see below) as between-subject factor. The ANOVAs reveal a strong effect of the expression type (*F*(1,60) = 188.82 respectively *F*(1,60) = 46.39;*p* < 0.00001), but no significant influence of the stimulus type (*F*(2,60) = 0.0 respectively *F*(2,60) = 0.01; *p* > 0.99). For both avatars we found a significant interaction (*F*(2,60) = 4.51; *p* = 0.015 respectively *F*(2,60) = 3.15; *p* = 0.049). This implies that the tuning to human expressions is narrower than that for monkey expressions, independent of the chosen avatar that was used to display the motion.

### Robustness of results against variations of expressive features

One may ask whether the previous observations are robust with respect to variations of the chosen stimuli. First, monkey facial movements include *species-specific features*, such as ear motion, that are not present in human expressions. Do the observed differences between the recognition of human and monkey expressions depend on these features? We investigated this question by repeating the original experiment with a new set of participants, using stimuli for which the ear region was occluded. Figures 2C and D depict the corresponding fitted discriminant functions, which are quite similar to the ones without occlusion, characterized again by a high similarity in shape between the human and monkey avatar (ratio of different vs. shared variance: *q* = 5.77%; only 12% of the categorization responses over the 25 points in morphing space were significantly different between the two avatar types; *p* = 0.02). Figure 3C shows that also the corresponding tuning functions *D*_M_ and *D*_H_ are very similar to the ones for the non-occluded stimuli, and the associated threshold values (Fig. 3D) are not significantly different (see above).

A second possible concern is that the chosen prototypical expressions might specify different amounts of expressive or salient low-level features, for example due to species differences in the motion or between the anatomies of the human and the monkey face. In order to rule out the influence of such differences, we repeated the experiment using a set of dynamic expressions (with non-occluded ears) that was equilibrated in terms of the average amount of optic flow and deformation information. This equilibration was based on a pilot experiment (see Supporting Information) demonstrating that the expressiveness of the stimuli was best predicted by the two-dimensional deformation flow of the underlying mesh. This deformation flow was manipulated by computing morphs between the original prototypical expression trajectories and ones of neutral facial expressions, exploiting the Bayesian generative model. Separate for the two avatar types, we determined morph levels that resulted in equal values of the deformation flow for all prototypes, where we tried to match the flow of the most expressive prototype (‘monkey fear’ for the monkey avatar, and ‘human anger’ for the human avatar). We repeated the experiment with motion morphs based on these equilibrated prototypes.

The resulting fitted discriminant functions (Figures 2E and 2F) are more symmetrical along the axes of the morphing space than the original stimuli. This is corroborated by the fact that an *Asymmetry Index* (AI) that measures the deviation from a perfect symmetry with respect to the *e* and *s* axis (see Supporting Information) is significantly reduced for the data from the experiment with equilibrated stimuli (*AI* = 0.656 vs. 0.486; *t*(21) = 2.81; *p* = 0.01). Again, we found very similar shapes of the discriminant functions for presentation on the human and the monkey avatar (ratio of different vs. shared variance: *q* = 11.6%; only 8% of the categorization responses over the points in morphing space were significantly different; Fig 3B). Most importantly, also for these equilibrated stimulus sets, we found a narrower tuning for the human than for the monkey dynamic expressions (Fig. 3C), consistent with the results of the ANOVA for the threshold points of the tuning functions *D*_M_(*s*) and *D*_H_(*s*) of the non-equilibrated stimuli. An analysis of the steepness of the fitted tuning functions at the threshold points (Fig. 3E) shows, in addition, that the equilibration removes the steepness difference between the monkey and the human expressions, which is apparent in the data from the non-equilibrated stimuli. This is confirmed by 2-way ANOVAs for the original motion stimuli and the ones with occluded ears, which show a (marginally) significant influence of the avatar type (human vs. monkey) (*F*(1,40) = 6.3; *p* = 0.0162 respectively *F*(1,40) = 3.33; *p* = 0.076), but not of the expression type (human vs. monkey motion) and no interactions (*F*(1,40) respectively *F*(1,39) < 0.01; *p* > 0.93). Contrasting with this result, the ANOVA for the stimuli with equilibrated motion does not show any significant effects, neither of the factor avatar type, nor of the expression type, nor an interaction (*F*(1,44) < 0.4; *p* > 0.53). The equilibration thus levels out the steepness difference of the category boundary between the human and the monkey avatar, but it does not affect that tuning for human expressions is more precise than the one for monkey expressions. The sharper tuning for own-species expressions is thus not just a side effect of differences in the amount of low-level salient features of the chosen prototypical motion patterns.

## Discussion

Due to the technical difficulties of an exact control of dynamics of facial expressions^20,21^, in particular of animals, the computational principles of the perceptual representation of dynamic facial expressions remain largely unknown. Exploiting advanced methods from computer animation with motion capture across species and machine-learning methods for motion interpolation, our study reveals fundamental insights about the perceptual encoding of dynamic facial expressions across primate species. At the same time, the developed technology lays the ground for physiological studies with highly-controlled stimuli on the neural encoding of such dynamic patterns^12,18,22,23^.

Our first key observation was that facial expressions of macaque monkeys were learned very quickly by human observers, always requiring less than 10 stimulus repetitions. This was the case even though monkey expressions are quite different from human expressions, so that naïve observers cannot interpret them spontaneously. This fast learning might be a consequence of the high similarity of the neuro-muscular control of facial movements in humans and macaques^15^, resulting in a high similarity of the structural properties of the expression dynamics that can be exploited by the visual system for fast learning.

Second and unexpectedly from shape-based accounts for dynamic expression recognition, we found that the categorization of dynamic facial expressions was only very weakly influenced by the basic shape of the face, as parameterized by the avatar type (human vs. monkey). Neither did we find strong differences between categorization responses between the two avatars, nor did we find a better perceptual representation of species-specific dynamic expressions that matched the species of the avatar. Facial expression dynamics is thus represented largely independently of the basic shape of the face. Yet, we found a clear and highly robust own-species advantage^24,25^ in terms of the accuracy of the tuning for expression dynamics: The tuning along the species axis of our motion style space was narrower for human than for monkey expressions. This remained even true for stimuli that eliminated species-specific features, or that were carefully balanced in terms of the amount of low-level information.

Both key results support our initial hypotheses: Perception can exploit the similarity of the structure of dynamic expressions across different primate species for fast learning. At the same time, and consistent with a co-evolution of the visual processing of dynamic facial expressions with their motor control, we found a largely independent encoding of facial expression dynamics from basic facial shape. Such independence seems also in-line with results from functional imaging studies that suggest a modular representation of different aspects of faces^26,27^. At the same time, this principle seems difficult to reconcile with popular (recurrent) neural network models that represent facial expressions in terms of sequences of learned key-shapes^4,28^. Since the shape differences between human and the monkey faces are much larger than the ones between the keyframes from the same expression, the observed spontaneous generalization to dynamic expressions to faces from a completely different species seems difficult to account for by such models. A separate encoding of facial dynamics from facial shape also explains why humans easily recognize expressions from comic characters that are not even primates. Concrete circuits for such shape-independent encoding of expression dynamics might be based on optic-flow analysis. Alternatively, such representations might be based on vectorized or on norm-referenced encoding, where face deformations are represented in terms of differences relative to a learned neutral reference pose of the face^29–31^. It seems an interesting theoretical question how deep neural architectures can be combined with such physiologically-motivated encoding principles. Our novel technology for the generation of photo-realistic, and however highly-controlled cross-species dynamic facial expressions enables electrophysiological studies that clarify the exact underlying neural mechanisms.

## Methods

### Human participants

In total, 58 human participants (32 female) participated in the psychophysical studies. The age range was from 21 to 53 years (mean: 26.9, standard deviation 5.11). All participants had no prior experience with macaque monkeys and normal or to-normal corrected vision. Participants gave written informed consent and were reimbursed by 10 EUR per hour for the experiment. In total, 21 participants (11 female) were taking part in the first experiment using stimuli based on the original motion capture data and the experiment with occlusion of the ears. 12 participants (8 female) took part in the experiment with equilibrated motion of the prototypes. In addition, 16 participants (8 female) took part in a Turing test control experiment (see below), and 9 (5 female) participants took part in a control experiment to identify features that influence perceived expressiveness of the stimuli. All psychophysical experiments were approved by the Ethics Board of the University Clinic Tübingen and consistent with the rules of the Declaration of Helsinki.

### Stimulus presentation

Subjects were presented the stimuli watching a computer screen at a distance of 70 cm in a dark room, using *Matlab^®^* and the *Psychotoolbox (3.0.15)* library for stimulus presentation^32,33^. Each stimulus was repeated at maximum three times before asking for the responses, but participants could skip after the first presentation if they were certain about their responses. Participants were first asked whether the perceived expression was rather from a human or a monkey, and whether it was rather the first or the second expression. Responses were given by key presses. Stimuli for the two different avatar types were presented in different blocks, with 10 repeated blocks per avatar type.

### Equilibration of stimuli for amount of motion / deformation

Stimuli were balanced for their amount of expressive low-level cues based on a control experiment that tested the relationship between different measures characterizing the amount of low-level cues and the rated expressivity of the stimulus for a set of morphs between the original prototypical facial movements and neutral expressions (see Supporting Information). Such morphs were generated by weighting the original expression with the morph level *λ* and the neutral expression with the weight (1 – *λ*) The most predictive measure for expressiveness was the two-dimensional *motion flow MF* of the vertex positions of the surface match, which could be computed easily from the animations (see Supplementary Information for details). Stimuli were equilibrated by matching, separately for the two avatar types, this measure to the value of the prototype motion that resulted in the largest flow. For this purpose, we fitted (separately for each avatar) the relationship between the morph level *λ* and the motion flow *MF* by a logistic function of the form:

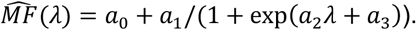

The inverse of this function was used to determine the values of the morph parameter *λ* that resulted in expressivities that matched the ones of the most expressive prototype motion.

### Statistical analysis

Statistical analyses were implemented using *Matlab^®^* and RStudio (3.6.2), using R and the package *lme4* for the mixed models of ANOVA.

Different GLMs for the modeling of the categorization data were fitted using the *Matlab Statistics Toolbox*. Models including different sets of predictors were compared using a step-wise regression approach. Models of different complexity were compared using the prediction accuracy and the Bayesian Information Criterion (BIC) as criteria.

Two statistical measures were applied in order to compare the similarity of the categorization responses for the two avatar types. First, we computed the ratio of the different vs. shared variance between the fitted discriminant functions, defined by the expression:

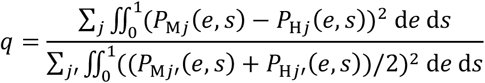

This ratio is zero if the discriminant functions for the human and the monkey avatar are identical. The *P*_M*j*_(*e*, *s*) and *P*_H*j*_(*e*, *s*) signify the fitted discriminant functions for the monkey and the human avatar with the category index *j*.

As second statistical analysis, we compared the multinomially distributed 4-class classification responses across the participants for the individual points in morphing space using a contingency table analysis that tested for significant differences between the two avatar types. Statistical differences were evaluated using a 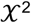-test, and for cases for which predicted frequencies were lower than 5, exploiting a bootstrapping approach^34^.

The species tuning functions *D*_H_(*s*) and *D*_M_(*s*) were fitted by the sigmoidal function *D_H,M_* = (tanh(ω(s — θ)) + 1)/2 with the parameter *θ* determining the threshold and *ω* the steepness. Differences of the tuning parameters *θ* were tested using 2-factor mixed-model ANOVAs (species-specific of motion (monkey vs. human) as within-subject factor, and experiment (original motion, occlusion of the ears, and equilibrated motion) as between-subject factor). Differences of the steepness parameters *ω* were tested using a within-subject two-factor ANOVAs.

## Acknowledgements

Special thanks to Tjeerd Dijkstra for the help with advanced statistical analysis techniques. We thank H. and I. Bülthoff for helpful comments. This work was supported by HFSP RGP0036/2016 and EC CogIMon H2020 ICT-23-2014/644727. MG was also supported by BMBF FKZ 01GQ1704 and BW-Stiftung NEU007/1 KONSENS-NHE. RS, SS, PD, and PT were supported by a grant from the DFG (TH 425/12-2), NVIDIA Corp.

## Author Contributions

MAG and PT developed the conceptual framework of the research. MAG, MS and NT designed the experiment. MS performed the experiment and did the statistical analysis. LS contributed to the experiment. NT, SS and PD recorded the motion capture data. RS cleaned, segmented and labeled motion data and provided advice about monkey communicative expressions. MAG, MS and NT wrote the initial version of the manuscript, and all authors interpreted the results and revised the manuscript.

