## Supplemental Information for "Shape-invariant perceptual encoding of dynamic facial expressions across species"

**This PDF file includes:**

Supplementary text
Figures S1 to S3
Tables S1 to S3
SI References

**Other supplementary materials for this manuscript include the following:**

Resource Table

### Supporting Information

#### Monkey subject

The monkey facial movements were recorded from a 9-year-old male rhesus monkey (Macaca mulatta), born in captivity and pair-housed. The monkey had previously been implanted with an individually adapted titanium head-post to allow head immobilization in unrelated neurophysiological experiments, and it had been trained to climb into a primate chair and to accept head fixation. The animal was in daily contact with other macaque monkeys and human caretaking personnel. The structural model of the monkey's head was derived from a T1-weighted MRI-scan with an isotropic resolution of 1mm. Motion capture recordings were compatible with the guidelines set by the National Institutes of Health and German national law and were approved by the local committee supervising the handling of experimental animals (Regierungspräsidium Tübingen, Abteilung Tierschutz, permit number N4/14). Human movements were recorded from a 40 years-old male human subject.

**Monkey avatar**

The highly realistic monkey avatar was generated exploiting state-of-the-art techniques from computer animation. Such techniques have been applied before for the realization of animation movies for cinemas[1]. However, monkey avatars of high quality have only very recently been developed for studies on static face perception[2], and to our knowledge this work is the first one exploiting motion capture from monkeys for generating such dynamic avatars. The head model was developed based on an MRI scan of an animal (Figure S1A). The scan provides a quite detailed model of the basic shape of the head, but is characterized by a highly irregular polygon structure, which makes it difficult to control the deformation during animation. In order to obtain a mesh model that can be manipulated more easily, we reduced the number of polygons and created a regularized mesh with clean edge loops (Figure S1B). This corrected mesh was adjusted for a neutral pose, and control points were specified that control the mesh deformation during animation. For the regularized mesh the weighting regions that determine the influences of the individual control points on the mesh could be exactly controlled. This developed ‘low-polygon’ model is useful for controlling the animation, but it lacks a lot of high-frequency details that are critical for a realistic appearance of the face. In order to add such details, we imported the model with the low polygon number into *Adobe Autodesk Mudbox*, a software that allows by subdivision to generate again a highly-regular mesh model with a high number of polygons. This model was further refined by a number of specific editing steps, including clay modeling in order to improve 3D shape details. Using a special tool (*Alpha Brushes*) additional texture details were added, such as wrinkles and pores (Figure S1C). To transfer the deformation from the low to the high-frequency polygon model we exported displacement maps in *Autodesk Mudbox,* which capture the differences between the low and high polygon-density models.

A particular challenge was the development of a realistic skin model that specifies believable color and reflectance properties. Skin surface textures were generated using photographs of a real monkey as a reference, and painting layer-wise color variations of the skin in order to approximate maximally its realistic appearance. Specifically, we used multiple layers of diffusive texture to model the translucent behavior of skin, separately for the deep layer, subdermal layer and epidermal layers (Figure S1D). For the deep layer, we hue-shifted the diffuse texture map towards red colors in order to model the deep vascularization while the color of the subdermal layers was shifted more towards yellow in order to simulate the fatty parts of the skin. In order to mimic the very thin superficial layer of dead skin we desaturated the diffuse texture for the epidermal layer. For realistic appearance it was also important to model the specularity of monkey skin, reproducing how light is reflected from the skin within different facial regions. For this purpose, we created two specular maps for the monkey's face, one simulating the basic specularity and one describing the oiliness vs. wetness of the skin (Figure S1E). Both material channels have an Index of Refraction (IOR) of 1.375, corresponding the IOR of water.

A final element that was essential for a realistic appearance was a realistic modeling of the fur. For this purpose, we exploited the built-in *XGen Interactive Groom* feature for hair creation of the animation software *Maya*. The overall appearance of the hair was controlled exploiting three control levels: The first level models the base of the fur, defining the direction of the hairs by control splines and adding some noise to model texture fluctuations and the matting of the fur. The second level models structures consisting of long hair, including the whiskers and the brows, using a smaller number of thick hairs. Believability and realism were increased further by adding a third layer of hair, also known as *Peach Fuzz* or *Vellus*, that consists of tiny hairs that are distributed within the face area. The final result is shown in Figure S1F.

**Human avatar**

The human avatar was based on a female face scan provided by the company *EISKO* (examples see Figure S2A). The commercial package includes also all main textures (diffuse map, specular map, base displacement map, etc.; Fig. S2B and S2C) for the neutral pose, as well as corresponding textures for face compression and stretching (Figures S2B and S2C). We applied just small color adjustments to the diffuse and specular map, similar to texture creation of the monkey head model. The *EISKO* model package included also a whole face rig with 154 blend shapes, suitable for changing of the face shape by blending (interpolation), resulting in naturally-looking shape variations (Figure S2A). The interpolation was driven by control points equivalent to the ones in the monkey model, defined by the motion-captured markers exploiting a ribbon-like structure that was inspired by the human muscle anatomy (Figure S2F). Using a tension map algorithm, we determined the local deformations of the texture from the mesh deformations relative to the neutral pose. For the generation of high-frequency details, contrasting with the approach for the monkey avatar, we employed a multichannel texture package from *TexturingXYZ.* This package provides diffuse maps and high-frequency details as displacement maps (pores, wrinkles, etc.) derived from a scanned real face. Exploiting the programs *R3dS Wrap* and *xNormal*, we transferred shape details similar to the ones of the monkey face to the human face model (Figure S2G). Hair animation used the same tools as for the monkey face. The final result is shown in Figure S2H.

### Motion capture

Motion capture was realized with a *VICON FX20* motion capture system with 6 cameras (focal length 24 mm) using a camera setting that was optimized for face capturing. We used 43 reflecting markers (2 mm) that were placed in the face, using a marker set that we developed ourselves (Figures 1B and S1B). Recording frequency was 120 Hz. Trajectories were preprocessed using *Nexus* 1.85 software by *VICON*, smoothed and segmented by an expert into individual facial expressions with a duration between 3 to 5 s. The trajectories were resampled with 150-time steps and 30 fps.

The monkey expressions were recorded from a 9-year-old male animal. The expressions were elicited by showing the animal different objects, including a screw driver, a mirror, and an unknown male human individual. The animal was head-fixed and observed the stimuli at a distance of 200 cm in front of the camera set-up. The recorded trajectories were segmented by a monkey expert, who had extensive experience with macaque monkeys on a daily basis.

The human marker set was corresponding to the one of the monkey, except that it lacked markers on the ears (Fig. S2D). The human actor was instructed to show two facial expressions 'anger' and 'fear'. Processing was identical to the marker trajectories of the monkey expressions.

### Motion morphing algorithm

In order to create continuous parameterized spaces of facial movements we exploited a motion morphing method that is based on a hierarchical Gaussian process model. The method is in principle real-time capable, thus allowing for instantaneous changes of motion style modulations based on on-line user input. This functionality was not critical for the experiments presented in this paper, but is used in ongoing experiments that build on the presented results.

Our motion morphing algorithm is based on a hierarchical probabilistic generative model that is learned from facial movement data. The architecture (Fig. S3A) comprises three layers. The lower two layers are formed by Gaussian process latent variable models (GP-LVMs)[3], and the highest layer is formed by a Gaussian Process Dynamical Model (GPDM)[4].

The facial motion was given by the *M*-dimensional trajectories of the control points, which were parameterized by an *N* x *D* time series matrix **Y** = $[\mathbf{y}_1, \ldots, \mathbf{y}_N]^{\mathrm{T}}$ with *N* = 600 (two expressions) or *N* = 900 (neutral expression included for equilibration) and *D* = 208 dimensions. The two GP-LVM layers reduce the dimensionality of the patterns in the high dimensional trajectory space in a nonlinear way. For this purpose, the first layer represents the trajectory points as nonlinear functions of a lower-dimensional hidden state variable, specifying the *N* x *M* matrix **H** = $[\mathbf{h}_1, \ldots, \mathbf{h}_N]^{\mathrm{T}}$. In our case the dimensionality *M* of the hidden variables $\mathbf{h}_n$ was six. Signifying by $(\mathbf{y}^d)^{\mathrm{T}}$ the row vectors of the matrix **Y**, the trajectory components are modelled in the form

$$\mathbf{y}^d = [f_1(\mathbf{h}_n), \ldots, f_1(\mathbf{h}_N)]^T + \boldsymbol{\varepsilon}_d \quad \text{with} \quad \boldsymbol{\varepsilon}_d \sim \boldsymbol{\mathcal{N}}(\boldsymbol{\varepsilon}_d|\mathbf{0}, \sigma^2\mathbf{I}),$$

where the variables $\boldsymbol{\varepsilon}_m$ specify independent Gaussian noise vectors, and with the function $f_1$ being drawn from a Gaussian process $f_1 \sim \mathcal{GP}(\mathbf{0}, k_1(\mathbf{h}, \mathbf{h}'))$, i.e. all vectors of the form $\mathbf{f_1} = [f_1(\mathbf{h}_n), \ldots, f_1(\mathbf{h}_N)]^T$ are distributed according to the Gaussian distribution $\boldsymbol{\mathcal{N}}(\mathbf{f}|\mathbf{0}, \mathbf{K})$ with the covariance matrix $\mathbf{K}$ whose elements are specified by a kernel function $k_1$ in the form $K_{nn'} = k_1(\mathbf{h}_n, \mathbf{h}_{n'}) + \gamma_1 \delta_{nn'}$. The kernel function is given by a linear combination of two types of kernels, a radial basis function kernel and a linear kernel:

$$k_1(\mathbf{h}, \mathbf{h}') = \gamma_2 \exp(-\beta_1 |\mathbf{h} - \mathbf{h}'|^2) \;\; + \gamma_3 \, \mathbf{h}^{\mathrm{T}} \mathbf{h}'$$

The RBF part allows to capture nonlinear structures in the data, while the linear kernel supports smooth and linear inter- and extrapolation in the pattern space. In addition, we found that the Kronecker delta part of the Kernel matrix is critical for the smoothness of the learned trajectories in the latent space. The parameter $\beta_1$ specifies the inverse width of the Gaussian radial basis functions.

The second layer of the model is defined exactly as the first layer. Here the dimensionality of the variables $\mathbf{h}_n$ is further reduced by generating by a nonlinear mapping from a hidden state variables $\mathbf{x}_n$ with *Q* = 2 dimensions, which defines the matrix **X** = $[\mathbf{x}_1, \ldots, \mathbf{x}_N]^{\mathrm{T}}$. Like in the first layer, the nonlinear mappings between the components of the variables **x** and **h** are defined by functions drawn from a Gaussian process $f_2 \sim \mathcal{GP}(\mathbf{0}, k_2(\mathbf{x}, \mathbf{s}, \mathbf{e};\, \mathbf{x}', \mathbf{s}', \mathbf{e}'))$, where the hyper-parameters of the kernel function differ from the ones of the kernel $k_1$. In addition, the kernel of this layer depends on the style vector variables **e** and **s.** These variables enter the kernel of the Gaussian process as multiplicative linear kernel terms:

$$k_2(\mathbf{x}, \mathbf{s}, \mathbf{e};\, \mathbf{x}', \mathbf{s}', \mathbf{e}') = \gamma_4 \, \mathbf{s}^{\mathrm{T}} \mathbf{s}' \;\; \mathbf{e}^{\mathrm{T}} \mathbf{e}' \exp(-\beta_2 |\mathbf{x} - \mathbf{x}'|^2) \;\; + \gamma_5 \, \mathbf{x}^{\mathrm{T}} \mathbf{x}' \,.$$

The random variables **e** and **s** encode the motion style using one-out-of-2 encoding, and they were estimated form the training data together with the state variables $\mathbf{x}_n$ using a maximum-likelihood approach. This parametrization turns out to be favorable to separate the different style components and the motion content in the latent space similar to multi-factor models[5]. We constrained the style vectors for all trials of the training data that represented the same motion style (e.g. ‘expression 1’, ‘human motion’) to be equal. In this way, the training data specify estimates $\hat{\mathbf{e}}_1$ and $\hat{\boldsymbol{e}}_2$ that correspond to averages of the expression types 1 and 2, and estimates $\hat{\mathbf{s}}_{\mathrm{M}}$ and $\hat{\mathbf{s}}_{\mathrm{H}}$ that correspond to the average monkey and the human expressions. In order to generate new intermediate motion styles, we ran the learned Gaussian model in a generative mode, fixing the values of these style vectors to blends between these estimates. The style vectors for the motion morphs as functions of the style parameters *e* and *s,* as discussed in the main part of the paper, were given by the relationships: $\boldsymbol{e} = e \cdot \hat{\boldsymbol{e}}_1 + (1-e) \cdot \hat{\boldsymbol{e}}_2$ and $\boldsymbol{s} = s \cdot \hat{\mathbf{s}}_{\mathrm{M}} + (1-s) \cdot \hat{\mathbf{s}}_{\mathrm{H}}$ respectively.

The highest level of the probabilistic model approximates the dynamics of the trajectories of the hidden state variables $\mathbf{x}_n$ using a nonlinear extension of an auto-regressive model, which is known as Gaussian Process Dynamical Model (GPDM). For this purpose, the state dynamics is modeled as function of the 2-dimensional hidden state variable $\mathbf{x}_n$ that obeys the nonlinear dynamics

$$\mathbf{x}_n = f_3\,(\mathbf{x}_{n-1}, \mathbf{x}_{n-2}) + \boldsymbol{\xi}_n \ ,$$

where $\boldsymbol{\xi}_n$ is isotropic white Gaussian noise, and where the nonlinear function $f_3$ is again specified by a Gaussian process. The hidden state dynamics can again be learned using a GP-LVM framework[4], where we used a kernel function of the form:

$$k_3(\mathbf{x}_{n-1}, \mathbf{x}_{n-2}; \mathbf{x}_{n'-1}, \mathbf{x}_{n'-2}) = \gamma_6 \exp(-\beta_3 |\mathbf{x}_{n-1} - \mathbf{x}_{n'-1}|^2 - \beta_4 |\mathbf{x}_{n-2} - \mathbf{x}_{n'-2}|^2) \\ + \gamma_7\, \mathbf{x}_{n-1}^{\mathrm{T}} \mathbf{x}_{n'-1} + \gamma_8 \mathbf{x}_{n-2}^{\mathrm{T}} \mathbf{x}_{n'-2}.$$

To determine the parameters of the GP-LVMs we maximized the logarithm of their posterior likelihood and fitted all hyper-parameters using a scaled conjugate gradient algorithm[6]. Since the evaluation of the posterior requires the inversion of a kernel matrix with a dimensionality that is given by all pairs of latent points, its direct implementation is computationally infeasible for large data sets. To render this inversion feasible we applied a sparse approximation method that approximates the posterior distribution based on a low number of inducing points in the hidden spaces (see[7]). The model was trained using six motion-captured example trajectories for each of two basic human and monkey expressions, sampled with 150-time steps. Training using an AMD Ryzen Threadripper 1950X CPU with 32 cores with a clock frequency of 3.4 GHz took about 1.5 hours. The most important parameters of the algorithm are summarized in Table S1.

### Turing test experiment

The described motion morphing algorithm interpolates between the original motion-captured movements in space-time. It was critical to verify that the morphing algorithm does not destroy the naturalness of the facial movements, at least for the prototypical expressions between which we blended. In order to verify this question, we realized a *Turing test* experiment that included 16 new participants. They had to discriminate between animations with original motion capture data ('original trajectories') and ones generated with movements that were generated by the morphing algorithm ('algorithm-generated trajectories'). The movements generated with the algorithm approximated the prototype movements (the style variables *e* and *s* being 0 or 1). In order to induce some variability, we used three different motion capture trials of each of the original human and monkey expressions, and their approximations based on the morphing model. The compared stimulus pairs were presented sequentially, and motions were presented in a block-randomized order 20 times, in separate blocks for the two avatar types. To verify that participants can pick up artifacts in the animations at all, we added a further condition where instead of movements generated by the morphing algorithm we used control movements, which were generated by reversing the temporal order of short 4-frame segments in the original motion-captured movements. Animations with these control movements also had to be distinguished from ones with the original motion capture data.

The results of this control experiments are shown in Figure S3B. The accuracy of the detection of original motion capture data as opposed to the generated one was 40.6% for the monkey avatar and 47.5% for the human avatar. Compared to the chance probability 0.5 both values are significantly lower ($\mathcal{X}^2(1,16) = 18.18;\ p < 0.001$ for the monkey and $\mathcal{X}^2(1,16) = 11.43;\ p < 0.001$ for the human avatar). This implies that the animations using motion capture data were judged even less frequently as 'original trajectories' than the animations generated with our motion synthesis algorithm. The morphing algorithm thus does not degrade the perceived naturalness of the motion. The even higher perceived naturalness of the algorithm-generated motion likely is a consequence of the motion being slightly more smooth, due to the smoothing properties of Gaussian process models. The artificial control movements were detected with very high reliability, as indicated by the high accuracies 96.88% for the monkey and 96.56% for the human avatars, which are highly significantly different from chance level ($\mathcal{X}^2(1,16) = 35.85;\ p < 0.001$ vs. $\mathcal{X}^2(1,16) = 34.93;\ p < 0.001$).

**Testing different low-level cues predicting expressivity**

Since we found for natural dynamic expressions that a larger part of the tested perceptual space was classified as monkey rather than as human expressions (cf. Figure 3C), we suspected this result to be a potential consequence of monkey expressions specifying more salient low-level features, e.g. local motion or geometrical deformations. In order to control for this variable, we created a second stimulus set for which the amount of low-level information was balanced. Since it was a priori unknown which type of low-level information drives the expressivity of facial expressions we tested a total of 9 possible measures, quantifying the amount of low-level features in a separate psychophysical experiment with 9 participants. These measures were: two-dimensional optic flow computed with a Horn-Schunck algorithm (*MATLAB®* implementation) from the movies, the absolute spatial deformation relative to the neutral frame, and the motion flow computed either from the control point trajectories or from the regularized mesh points, either in three dimensions or after projection to the two-dimensional image plain. The *spatial deformation* relative to the neutral frame was quantified using the measures:

$$DF = \sum_{t=1}^{N} \|\mathbf{X}_t - \mathbf{X}_0\|_2 \ ,$$

where $\mathbf{X}_t$ signifies a vector that contains the relevant control point or (two- or three-dimensional) mesh point coordinates, and where $N$ is the number of stimulus frames. Likewise, the *motion flow* was defined by the quantity:

$$MF = \sum_{t=1}^{N-1} \|\mathbf{X}_t - \mathbf{X}_{t-1}\|_2 \ .$$

For the true optic flow, the motion measure was computed by summing up the absolute values of all estimated local motion vectors across the image. The stimuli for this experiment were motion morphs between each of the four prototype expressions (two human and two monkey expressions) and a neutral expression. The original expression entered the motion morph with a weight of $\lambda$, and the neutral expression with the weight of $(1-\lambda)$ where the morphing weight was adjusted to obtain the same amount of low-level information in all adjusted prototypes.

In order to cut down the number of measures for the amount of low-level information in the first place, we generated a set of face motion stimuli with reduced and exaggerated expressivity, separately for the two face avatars, by choosing 6 different values for the morphing weight $\lambda$ (values 0 – 25 – 50 – 75 – 100 – 125% for the monkey expressions, and the values 0 - 37.5 – 75 - 112.5 – 150% for the human expressions). For all rendered movies we computed the 9 different measures for the low-level feature content and analyzed their dependence on the morphing weight $\lambda$ and their similarities. We found that measures *DF* and *MF* computed from the two-dimensional and three-dimensional mesh coordinates, and the control points were very highly correlated ($r > 86.24$, $r_{average} = 98.74$; $p < 0.0403$). The mesh point-based measures were monotonically increasing functions of the morphing level $\lambda$. This was not the case for the quantities computed from the control point trajectories, so that we discarded the measures derived from the control point trajectories from the balancing of the stimuli. Because of the high correlation between the measures computed from the two- and three-dimensional mesh-point trajectories, and the higher similarity of the two-dimensional trajectories with image motion we kept only the measures computed form the two-dimensional mesh-point trajectories for the further analysis. In addition, we tested the optic flow computed by the optic flow algorithm from pixel images as a third possible predictor of the low-level information. For each of these three predictors, we constructed a balanced stimulus set by adjusting the morph levels of all prototypes, except for the one with the lowest low-level feature content, in order to match their low-level information contents. As result, we obtained three balanced sets of stimuli, each with four dynamic expressions, separately for each avatar type.

All stimuli were shown in block-wise randomized order to the participants who had to rate their expressivity on a 9-point Likert scale. Figure S3C shows the averages and variabilities across subjects of the perceived expressivity.

For the human avatar, the stimuli whose expressivity was balanced using the motion flow measure *MF* showed the smallest variability across participants, and the largest expressivity. For the monkey avatar, the expressivity was rated similarly for stimuli balanced using the measures *MF* and *DF*, while it was significantly lower for stimuli balanced using the optic flow ($t(275) = 2.8$; $p = 0.0054$ and $t(269) = 3.95$; $p < 0.001$). A step-wise regression analysis, where we predicted the expressivity ratings from the remaining measures (*MF* and *DF* computed from the two-dimensional mesh motion). showed that the motion flow *MF* is sufficient, while the other predictor *DF* did not add significant additional information. Using a model comparison analysis exploiting the Bayesian Information Criterion (BIC), we found no significant difference of the explanatory values of the models including the predictor *MF*, and the predictors *MF* and *DF*: $\mathcal{X}^2(1{,}284) = 3.49; p = 0.062$.

**Asymmetry index**

The deviation of the four discriminant functions $P_i(e, s)$ from the completely symmetrical case, where all four discriminant functions have the same basic shape (with their peaks centered on the different prototypes) was quantified by defining and asymmetry index AI. This index is exactly zero if the four discriminant functions are exactly symmetrical with respect to the axes $e = 0.5$ and $s = 0.5$. This implies the symmetry relationship: $P_1(e, s) = P_2(1 - e, s) = P_3(e, 1 - s) = P_4(1 - e, 1 - s)$. In order to compute the index, we first computed a symmetrized average of all four discriminant functions according to the formula:

$$P_{\text{sym}}(e, s) = \frac{P_1(e, s) + P_2(1 - e, s) + P_3(e, 1 - s) + P_4(1 - e, 1 - s)}{4}$$

Likewise, we defined a standard deviation relative to this symmetrized average by the expression $\text{SD}_{\text{sym}}(e, s) = \sqrt{\text{Q}_{\text{sym}}(e, s)/3}$ with the least square deviation sum:

$$\text{Q}_{\text{sym}}(e, s) = (P_1(e, s) - P_{\text{sym}}(e, s))^2 + (P_2(1 - e, s) - P_{\text{sym}}(e, s))^2 +$$

$$(P_3(e, 1 - s) - P_{\text{sym}}(e, s))^2 + (P_4(1 - e, 1 - s) - P_{\text{sym}}(e, s))^2$$

The asymmetry index was defined by the expression:

$$\text{AI} = \frac{\iint_0^1 \text{SD}_{\text{sym}}(e, s)\,\text{d}e\,\text{d}s}{\iint_0^1 P_{\text{sym}}(e, s)\,\text{d}e\,\text{d}s}$$

The asymmetry index increases with the deviation from the completely symmetric case, where all four categories are represented equally well.

**Figure S1**

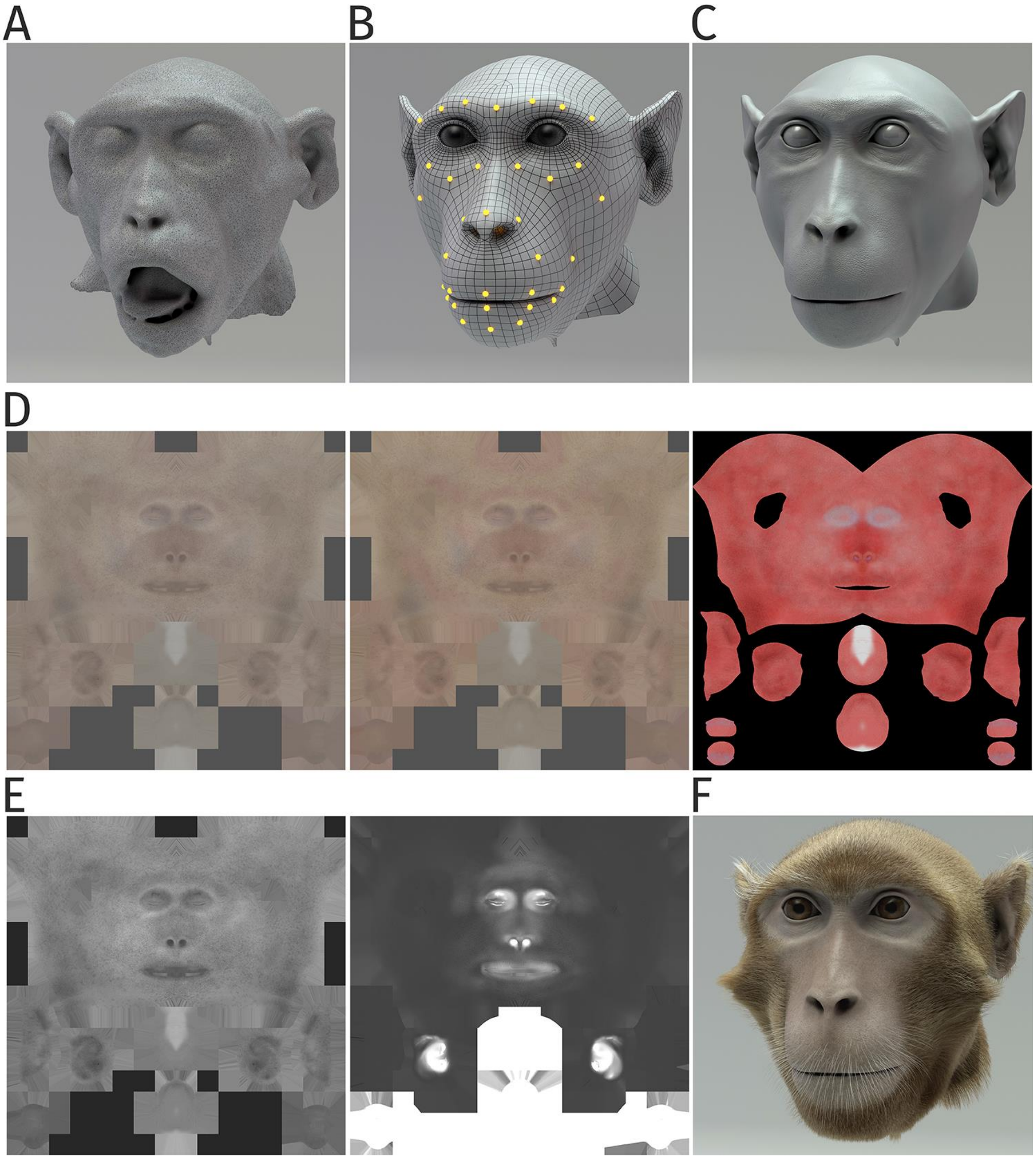


**Figure S1. Details of generation of the monkey head model.** (A) Irregular surface mesh resulting from a magnetic resonance scan of a monkey head (B) Face mesh whose deformation is following control points specified by motion-captured markers. (C) Surface with high polygon number derived from the mesh in (B), applying displacement texture maps, including high-frequency details such as pores and wrinkles. (D) Skin texture maps modeling the epidermal layer (left), the dermal layer (middle), and the subdermal layer (right panel). (E) Specularity textures modeling the reflection properties of the skin; overall specularity (left) and map specifying oily vs. wet appearance (right panel). (F) Complete monkey face model including the modeling of fur and whiskers.

**Figure S2**

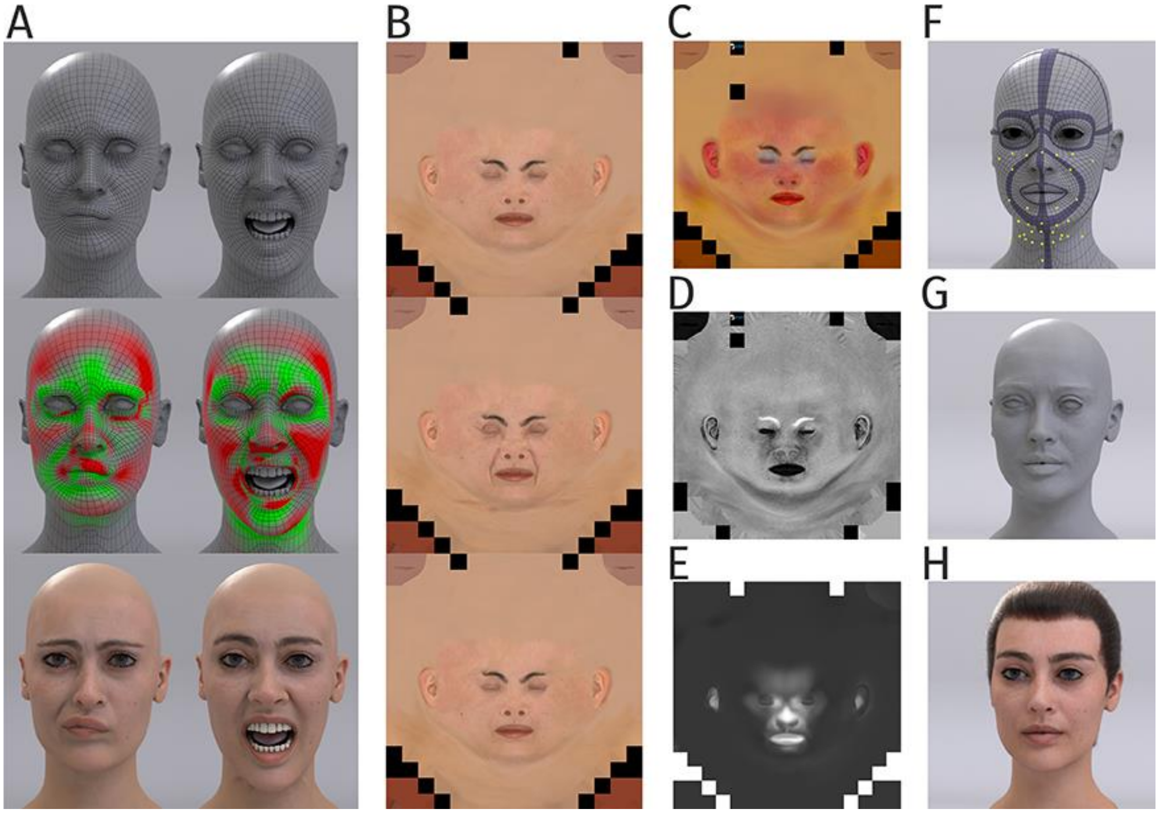


**Figure S2. Details of generation the human head model.** (A) Human face mesh and deformations by a blendshape approach, where face poses are driven by the 43 control points (top panel). Tension map algorithm computes compression (green) and stretching (red) of mesh during animation (middle panel). Corresponding texture maps were blended (bottom panel). (B) Examples of diffuse texture maps (top panel) with additional maps for stretching (middle panel) and compression (bottom panel). (C) Subsurface color map modeling the color variations by light scattering and reflection by the skin. (D) Specular map modeling the specularity of the skin. (E) Wetness map modeling the degree of wetness vs. oilynes od the skin. (F) Regularized basic mesh with embedded muscle-like ribbon structures (violet) for animation. Yellow points indicate the control points defined by the motion capture markers. (G) Mesh with additional high-frequency details. (H) Final human avatar including hair animation.

**Figure S3**

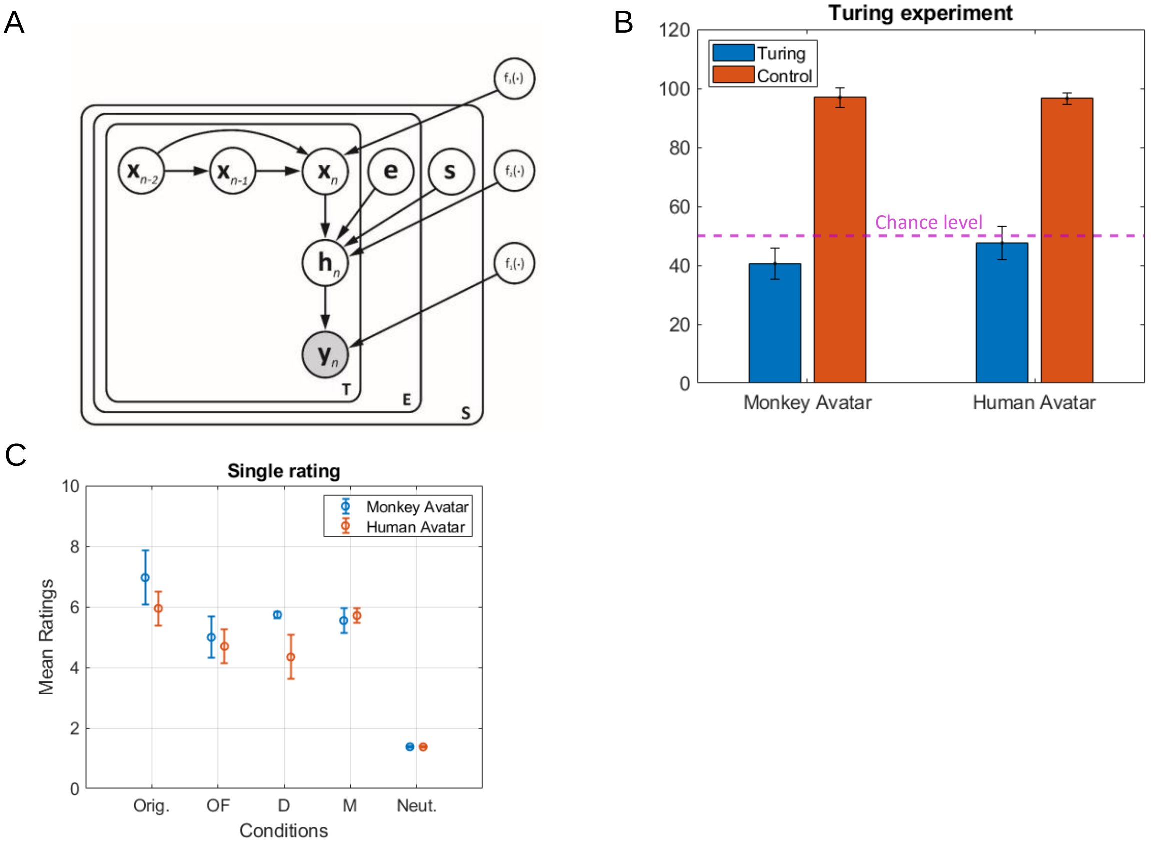


**Figure S3. Motion morphing algorithm and additional results.** (A) Graphical model showing the generative model underlying our motion morphing technique. The hierarchical Bayesian model has three layers, reducing subsequently the dimensionality of the motion data $\mathbf{y}_n$. The top layer models the trajectory in latent space using a GPDM. The vectors **e** and **s** are additional style vectors encode the expression type and the species type. They are binomially distributed. Plate notation indicates the replication of model components for the encoding of the temporal sequence, and the different styles. Nonlinear functions are realized as samples form Gaussian processes with appropriately chosen kernels. (Details see text.) (B) Results from Turing test experiment. Accuracy for the distinction between animations with original motion capture data and trajectories generated by our motion morphing algorithm is close to chance level (dashed line), opposed to the accuracy for the detection of small motion artifacts in control stimuli, which was almost one for both avatar types. (C) Expressiveness ratings for stimulus sets with and adjustment of expressiveness using different types of low-level predictors, separately for the monkey and the human avatar. (Conditions: Original stimuli; OF: optic flow; 2D metrics *D* or *M* (see text); Neutral: neutral facial expression with minimal motion). Error bars indicate standard deviations over participants.

**Table S1**

| Parameters of motion morphing algorithm | | |
|---|---|---|
| **Parameters** | **Description** | **Value** |
| $D$ | Data dimension | 208 |
| $M$ | First layer dimension | 6 |
| $Q$ | Second layer dimension | 2 |
| $T$ | Number of samples per trial | 150 |
| $S$ | Number of species | 2 |
| $E$ | Number of expressions | 2 or 3 |
| $N$ | Number of all samples | *T * S * E* |
| **Hyper Parameters** (learned) | | **Size** |
| $\beta_1$ | Inverse width of kernel $k_1$ | 1 |
| $\beta_2$ | Inverse width of kernel $k_2$ | 1 |
| $\beta_3$ | Inverse width for non-linear part one of kernel $k_3$ | 1 |
| $\beta_4$ | Inverse width for non-linear part two of kernel $k_3$ | 1 |
| $\gamma_1$ | Precision absorbed from noise term $\boldsymbol{\varepsilon}_d$ | 1 |
| $\gamma_2$ | Variance for non-linear part of $k_1$ | 1 |
| $\gamma_3$ | Variance for linear part of $k_1$ | 1 |
| $\gamma_4$ | Variance for non-linear part of $k_2$ | 1 |
| $\gamma_5$ | Variance for linear part of $k_2$ | 1 |
| $\gamma_6$ | Variance for non-linear part of $k_3$ | 1 |
| $\gamma_7$ | Variance for linear part one of $k_3$ | 1 |
| $\gamma_8$ | Variance for linear part two of $k_3$ | 1 |
| **Variables** | | **Size** |
| $\mathbf{Y}$ | Data | *N* x *D* |
| $\mathbf{H}$ | Latent variable of first layer | *N* x *M* |
| $\mathbf{X}$ | Latent variable of second layer | *N* x *Q* |
| $\hat{\mathbf{s}}_{\mathrm{M}}$ | Style variable vector for monkey species | *S* x *1* |
| $\hat{\mathbf{s}}_{\mathrm{H}}$ | Style variable vector for human species | *S* x *1* |
| $\hat{\mathbf{e}}_{\mathbf{1}}$ | Style variable vector for expression one | *E* x *1* |
| $\hat{\mathbf{e}}_{\mathbf{2}}$ | Style variable vector for expression two | *E* x *1* |

**Table S1. Parameters of the Bayesian motion morphing algorithm.** The observation matrix **Y** is formed by *N* samples of dimension *D*, where *N* results from *S* * *E* trails with *T* time steps. The dimensions *M* and *Q* of the latent variables were manually chosen. The integers *S* and *E* specify the number of species and expressions (two in our case).

**Table S2**

| ANOVAS | | | | | | |
|---|---|---|---|---|---|---|
| **Threshold** | *Monkey Avatar* | **Sum of Square** | **df** | **Mean Square** | **F** | **p** |
| | Stimulus type | 0,00 | 2 | 0,00 | 0,00 | 0,999 |
| | Expression type | 1,20 | 1 | 1,20 | 188,83 | 0,000 |
| | Stimulus * Expression | 0,06 | 2 | 0,03 | 4,51 | 0,015 |
| | Error | 0,42 | 60 | 0,01 | | |
| | Total | 1,72 | 65 | | | |
| | *Human Avatar* | | | | | |
| | Stimulus type | 0,00 | 2 | 0,00 | 0,01 | 0,993 |
| | Expression type | 0,40 | 1 | 0,40 | 46,37 | 0,000 |
| | Stimulus * Expression | 0,05 | 2 | 0,03 | 3,15 | 0,049 |
| | Error | 0,57 | 60 | 0,01 | | |
| | Total | 1,02 | 65 | | | |
| **Steepness** | *Original Motion Stimulus* | | | | | |
| | Avatar type | 376,68 | 1 | 376,68 | 6,3 | 0,016 |
| | Expression type | 0,36 | 1 | 0,36 | 0,01 | 0,939 |
| | Avatar * Expression | 0,16 | 1 | 0,16 | 0 | 0,959 |
| | Error | 2391,21 | 40 | 59,78 | | |
| | Total | 2768,41 | 43 | | | |
| | *Occluded Motion Stimulus* | | | | | |
| | Avatar type | 286,17 | 1 | 286,17 | 3,33 | 0,076 |
| | Expression type | 0,02 | 1 | 0,02 | 0 | 0,988 |
| | Avatar * Expression | 0,00 | 1 | 0,00 | 0 | 0,995 |
| | Error | 3094,54 | 36 | 85,96 | | |
| | Total | 3380,73 | 39 | | | |
| | *Equilibrated Motion Stimulus* | | | | | |
| | Avatar type | 1,57 | 1 | 1,57 | 0,4 | 0,533 |
| | Expression type | 0,25 | 1 | 0,25 | 0,06 | 0,803 |
| | Avatar * Expression | 0,02 | 1 | 0,02 | 0 | 0,945 |
| | Error | 174,76 | 44 | 3,97 | | |
| | Total | 176,60 | 47 | | | |

**Table S2. Detailed results of the 2-way ANOVAs.** ANOVA for the Threshold: 2-way mixed model with expression type as within-subject factor, and the stimulus type as between-subject factor for both, Monkey and Human Avatar. Steepness: 2-way ANOVA with Avatar type and Expression factor for each stimulus motion type (Original, Occluded and Equilibrated). The mean Square is define as: Mean Square = Sum of Square / df, with df = degree of freedom. In bold: Analysis Condition or Column Title. In italic: sub category of experiment design.

### Data and code availability

The pre-processed motion capture data that was used to train our Bayesian algorithm and the stimuli used in all the experiments are available on GitLab (https://hih-git.neurologie.uni-tuebingen.de/ntaubert/FacialExpressions). Data of all participants together with the *Matlab*® code and *R* scripts to run and analyze the experiments are also available at the same location. A Readme file guides the reader how to recreate all experiments and reproduce all figures.

### Resource Table

| RESOURCE | | |
|---|---|---|
| **Software and Algorithms** | **SOURCE** | **IDENTIFIER** |
| Custom-written software written in C# | This study | https://hih-git.neurologie.uni-tuebingen.de/ntaubert/FacialExpressions |
| C3Dserver | Website | https://www.c3dserver.com |
| Visual C++ Redistributable for Visual Studio 2012 Update 4 x86 and x64 | Website | https://www.microsoft.com/en-US/download/details.aspx?id=30679 |
| AssimpNet | Website | https://www.nuget.org/packages/AssimpNet |
| Autodesk Maya 2018 | Website | https://www.autodesk.com/education/free-software/maya |
| Matlab 2019b | Website | https://www.mathworks.com/products/matlab.html |
| Psychophysics toolbox 3.0.15 | Website | http://psychtoolbox.org/ |
| R 3.6 | Website | https://www.r-project.org/ |
| **Deposited Data** | | |
| Training data for interpolation algorithm | This study | https://hih-git.neurologie.uni-tuebingen.de/ntaubert/FacialExpressions/tree/master/Data/MonkeyHumanFaceExpression |
| Stimuli for experiments | This study | https://hih-git.neurologie.uni-tuebingen.de/ntaubert/FacialExpressions/tree/master/Stimuli |
